# Parameter-free clear image deconvolution (CLID) technique for single-frame live-cell super-resolution imaging

**DOI:** 10.1101/2024.09.11.612552

**Authors:** Fudong Xue, Wenting He, Zuo’ang Xiang, Jun Ren, Chunyan Shan, Lin Yuan, Pingyong Xu

**Author notes:** These authors contributed equally to this work (Fudong Xue, Wenting He).

## Abstract

Advancing single-frame imaging techniques beyond the diffraction limit and upgrading traditional wide-field or confocal microscopes to super-resolution (SR) capabilities are greatly sought after by biologists. While enhancing image resolution by deconvolving noise-free images is beneficial, achieving a noise-free image that maintains the distribution of signal intensity poses a challenge. We first developed a denoising method utilizing reversibly switchable fluorescent proteins through <u>s</u>ynchronized <u>s</u>ignal <u>s</u>witching (3S). Additionally, we introduced a denoising neural network technique, 3Snet, which combines supervised and self-supervised learning using 3S denoised images as the ground truth. These approaches effectively eliminate noise while maintaining fluorescence signal distribution across camera pixels. We then implemented <u>cl</u>ear image <u>d</u>econvolution (CLID) on both 3S and 3Snet denoised images to develop SR techniques, named 3S-CLID and 3Snet-CLID. Notably, 3Snet-CLID boosts the resolution of single fluorescence images from wide-field and spinning-disk confocal microscopies by up to 3.9 times, achieving a spatial resolution of 65 nm, the highest in such imaging scenarios without an additional SR module and complex parameter setting. 3Snet-CLID enables dual-color single-frame live-cell imaging of various subcellular structures labeled with conventional fluorescent proteins and/or dyes, allowing observations of dynamic cellular processes. We expect that these advancements will drive innovation and uncover new insights in biology.

---

Imaging the dynamic subcellular structures of living cells requires high temporal and spatial resolution to precisely capture positioning while minimizing motion artifacts. While traditional wide-field (WF) and confocal microscopy provide single-frame temporal resolution and simplicity, their spatial resolution is restricted by the diffraction limit. In contrast, modern super-resolution (SR) microscopy techniques, including structured illumination microscopy (SIM)^1^, stimulated emission depletion (STED)^2^ microscopy, photoactivated localization microscopy (PALM)^3^, stochastic optical reconstruction microscopy (STORM)^4^, and optical fluctuation imaging (SOFI)^5^, have revolutionized the ability to visualize cellular structures and molecular interactions at the nanoscale. However, these technologies often require costly and complex equipment or sacrifice temporal resolution to achieve heightened spatial resolution using multiple image frames. There is a pressing need to push single-frame imaging capabilities beyond the diffraction limit and upgrade traditional WF or confocal microscopy to achieve SR capabilities.

Recently, deep learning-based SR (DLSR) methods have advanced single-frame temporal resolution in WF microscopies by learning the end-to-end image transformation relationship^6–10^. However, training DLSR models requires a considerable amount of high-quality ground truth SR images, which is labor-intensive and often impractical due to the rapid dynamics in biological specimens^8, 10, 11^. Additionally, integrating expensive and complex SR equipment into a WF or confocal system is necessary if SIM or STED data are used as the ground truth. The SR reconstruction and denoising is challenging as there are multiple possible mapping relationships between the low-resolution and SR data, making it an ill-posed problem^7^. This renders the DLSR method structure-dependent, meaning that a deep learning network trained on one organelle may not be effective for imaging a different organelle, potentially leading to artifacts. To address this issue, researchers have proposed various strategies^12–14^, such as developing more generalizable deep learning models, expanding training datasets to increase diversity and complexity, and integrating multimodal imaging technologies. While these approaches have improved the generalization and robustness of neural networks to some extent, they also introduce additional complexity, requiring more sophisticated imaging systems and larger datasets for training.

Deconvolution algorithms and other analytical model-based methods show promise in achieving single-frame SR imaging^9, 15–18^. For example, the iterative Richardson-Lucy (RL) deconvolution^19, 20^ has demonstrated improved resolution in synthetic, noise-free fluorescence images but struggles with real microscopic images due to noise dominance after a few iterations^18,21^. A more recent development is sparse deconvolution SR technology, which aims to reduce noise influence by incorporating sparsity and continuity as priori knowledges into the algorithm^18^. This approach has successfully enhanced the resolution of single-frame WF microscopy to 120-150 nm^18^. However, both sparse deconvolution and other analytical models rely on assumptions about specimen and image properties to enhance resolution^18, 22, 23^, with multiple parameters that require manual tuning, making the process time-consuming and heavily reliant on the selected parameter sets^18–20, 22^. While there are additional computational methods like MSSR^24^, A-PoD^25^, and ZS-DeconvNet^26^, that can achieve single-frame SR imaging, they often entail long computation times, exhibit a bias towards structural imaging effects, and offer limited enhancements in WF or spinning-disk (SD) resolution.

In summary, there is a need for novel approaches to achieve single-frame SR imaging that can update commonly used equipment like WF and confocal microscopes without being reliant on specific subcellular structures. In comparison to the sparse method that use prior knowledge to constrain deconvolution amid noise presence, our study introduces a new technique called <u>cl</u>ear image <u>d</u>econvolution (CLID) to enhance resolution. The process involves utilizing a <u>s</u>ynchronized <u>s</u>ignal <u>s</u>witching (3S) denoising method, which leverages reversibly switchable fluorescent proteins (RSFPs) to drastically improve the signal-to-noise ratio (SNR) of the image while maintaining the intensity distribution of the signal on pixels. The denoised clear image is then subjected to deconvolution (3S-CLID) to successfully achieve large field-of-view SR imaging of fixed cells. To enhance temporal resolution and realize single-frame SR imaging of live cells, we introduce a jointly supervised and self-supervised denoising deep learning network (3Snet) using the denoised image of the fixed cell with 3S as the ground truth. The deconvolution of the 3Snet-denoised clear image is defined as 3Snet-CLID. 3Snet-CLID can effectively denoise different cellular structures and enhance spatial resolution without relying on deep learning SR structures. Notably, it enables the achievement of a 65-nm resolution for conventional FP/dye-labeled samples on WF and SD confocal microscopy, significantly improving image contrast.

## Results

### Conception of 3S-CLID

While aiming for a completely noise-free image may be unrealistic, the simulated data^18^ convincingly illustrate the resolution improvement achieved by directly RL deconvolving noise-free images. This serves as a compelling motivation to progress in developing techniques for acquiring nearly noise-free images. We have named the method that applies direct RL deconvolution on a clear image closely representing the true signal to enhance resolution as clear image deconvolution (CLID). To achieve CLID, it is crucial to preserve signal integrity and distribution while minimizing noise. However, removing noise is challenging due to its various forms, such as Gaussian and Poisson noise. Current denoising techniques frequently sacrifice signal fidelity and distort pixel-wise distribution, making them unsuitable for direct RL deconvolution. To address this issue, we propose leveraging the light-modulated optical switching properties of RSFPs to differentiate between signals and noise. We developed a denoising strategy depicted in Fig. 1a, involving two steps: signal acquisition and noise removal. Initially, Skylan-S (a green RSFP) underwent repeated switching cycles using 405 nm and 488 nm illumination for ON and OFF states, respectively. Multiple frames of ON and OFF images were captured under the same 488 nm illumination. We then conducted noise reduction by averaging multiple frames for both ON and OFF images to eliminate random Gaussian noise and signal-related Poisson noise. Subsequently, we subtracted the averaged OFF image (AVG(OFF)) from the averaged-ON image (AVG(ON)) to eliminate systematic noise, resulting in the final denoised image, referred to the clear image. This denoising approach was named <u>s</u>ynchronized <u>s</u>ignal <u>s</u>witching (3S) denoising. To further overcome the Nyquist sampling limit, we upscaled the denoised clear image by a factor of 2 using Lanczos^27^ (Extended Data Fig. 1) and applied deconvolution (3S-CLID) to enhance the resolution.

**Fig. 1.**
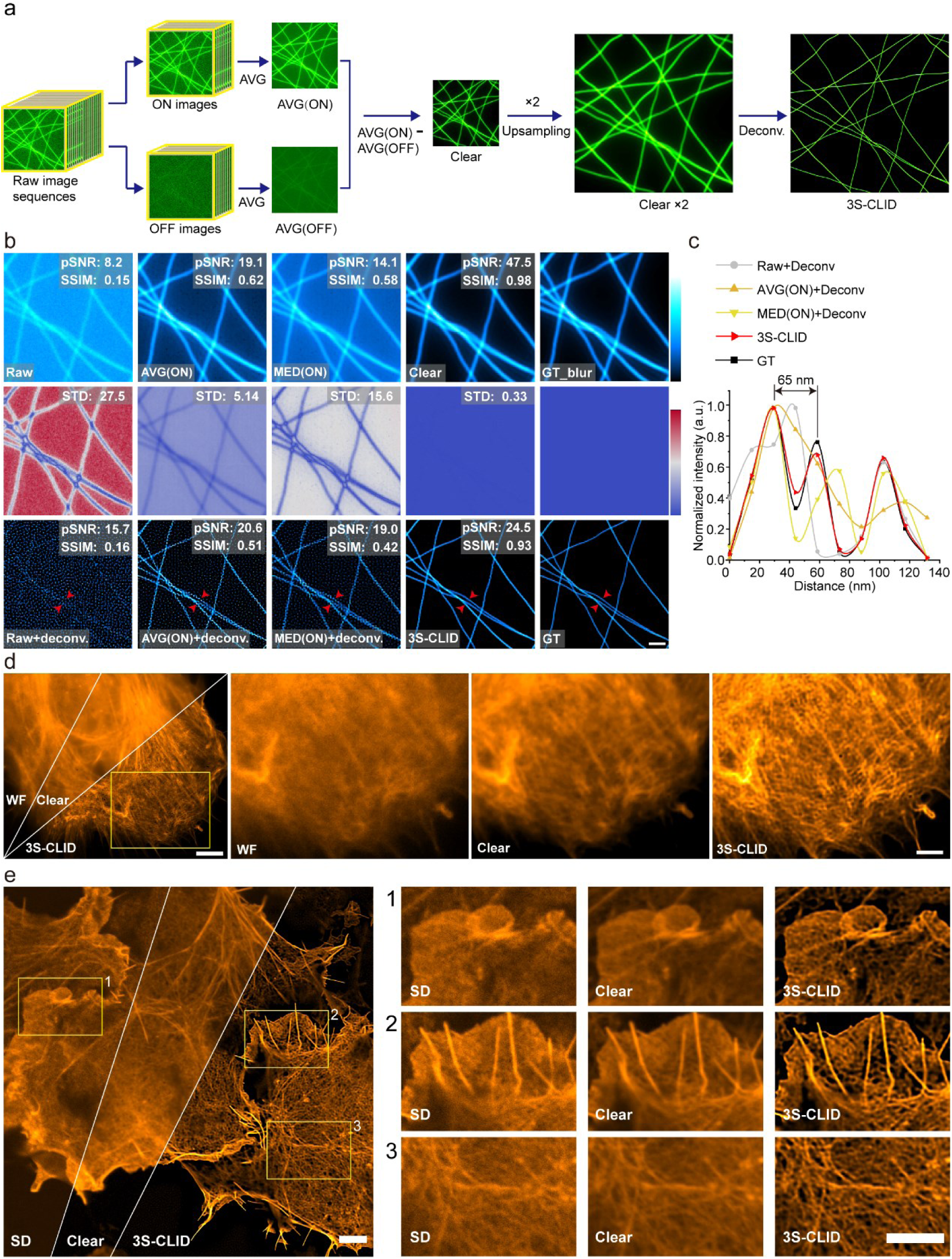
Workflow and validation of 3S-CLID on synthetic microtubule and cellular structure. **a**, Overview of 3S-CLID. The workflow involves 3S-denoising, where the image averaging of ON and OFF images, produced by RSFP optical switching, eliminate noise, and the averaged OFF image is subtracted from the averaged ON image to produce a denoised clear image. The denoised image is then upsampled and deconvolved using Lanczos and RL deconvolution to enhance resolution (**Methods**). **b**, Validation of 3S-CLID using synthetic microtubule. The synthetic structures were convoluted with a 250 nm PSF as blured ground truth. The noisy images were created by further injection of 5% Poisson noise and 5% Gaussian noise (**Methods**). Top row: peak signal-to-noise ratio (pSNR) and structural similarity (SSIM), from left to right: noisy image, image averaging of ON image sequences, median of ON image sequences, 3S-denoising result and blured ground truth. The color bar black to cyan means the relative intensity of images. Middle row: Data uncertainty results of Top row indicated by the averaged standard deviation (STD). The color bar blue to red represents the STD value. Bottom row: deconvolution of noisy image, deconvolution of averaged ON image, deconvolution of median averaged ON image, 3S-CLID result and ground truth. **c**, Normilized intensity profiles along the red arrow heads in the left ground truth (GT) and deconvoled images. **d**, 3S-denoised clear and 3S-CLID images of Lifeact-SkylanS-labeled F-actin in fixed U-2 OS cells under wide field microscopy and zoomed views from the yellow box in the left. **e**, 3S-denoised clear and 3S-CLID images of Lifeact-SkylanS-labeled F-actin in fixed U-2 OS cells imaged by Spining-disk (SD) confocal. Scale bars: 1 μm (**b**), 5 μm (**d**(left)**, e**(left)), 2 μm (**d**(right), **e**(right)).

### Evaluation of 3S-CLID on synthetic microtubule and cellular structure

We first evaluated the efficacy of 3S denoising using synthetic microtubules. To simulate real imaging conditions, images of a blank area without a sample were captured under WF illumination, and Gaussian and Poisson noise were added to the synthetic microtubule image as noise. The pixel size was set to 65 nm, consistent with the Flash 4.0 camera under a 100× objective. The results demonstrated that the clear image produced by the 3S denoising method significantly outperformed the images generated by image averaging (AVG(ON)) and pixel-wise median of image sequences (MED(ON)), with notably higher peak signal-to-noise ratio (pSNR) and structural similarity index (SSIM) (Fig. 1b). This indicates that 3S denoising generates higher quality image with less noise, portraying enhanced signal integrity and distribution compared to the commonly used image averaging method. In contrast, RL deconvolution of raw, AVG(ON) and MED(ON) images failed to separate 65 nm lines and generated artifacts, while 3S denoising followed by CLID achieved 65 nm resolution and enhanced image contrast (Fig. 1b and Fig.1c).

We proceeded by labeling cytoskeleton actin with RSFP and imaging fixed cells using WF and SD microscopy. The fluorescence signal acquisition was conducted following the procedure outlined in Fig. 1a. In both instances (Fig. 1d, WF and Fig. 1e, SD), 3S denoising significantly enhanced the SNR and structural continuity. Furthermore, 3S-CLID improved resolution and contrast, resolving finer structural details of actin bundles and microfilaments (Fig. 1d and Fig. 1e).

### Conception of 3Snet-CLID

Expanding on the achievements of 3S-denoising and 3S-CLID, we explored improving single-frame image quality by integrating 3S-CLID with specialized denoising networks designed to boost spatial resolution. In recent years, deep learning-based image denoising has emerged as a predominant method due to its remarkable capabilities^28^. While supervised deep learning denoising networks have exhibited superior performance, acquiring high-quality ground truth images for training poses challenges. Conventional denoising methods like utilizing high SNR images or filtering techniques often retain noise or introduce signal loss, resulting in suboptimal outcomes that are unsuitable for direct deconvolution to boost resolution. We postulate that the 3S denoising method, proficient in noise removal while faithfully preserving signal fidelity and pixel-wise distribution, can generate images serving as dependable and precise ground truth for training denoising neural networks. Inspired by the efficacy of supervised-to-self-supervised transfer learning, surpassing both supervised and self-supervised learning individually in denoising tasks^29^, we designed a deep learning denoising network based on U-net (termed 3Snet). This network integrates self-supervised and supervised denoising neural networks (Fig. 2a). By averaging ON images with varying frames, we generated noisy images with different noise levels, and randomly selected two noisy images at each noise level to train the self-supervised network for every training iteration. Concurrently, 3S-denoised images were utilized as ground truth for training supervision (Extended Data Fig. 2). The 3Snet network was employed to denoise single-frame images via WF or SD, subsequently upsampling and applying CLID to accomplish single-frame SR imaging (Fig. 2b). This methodology was termed 3Snet-CLID SR technology.

**Fig. 2.**
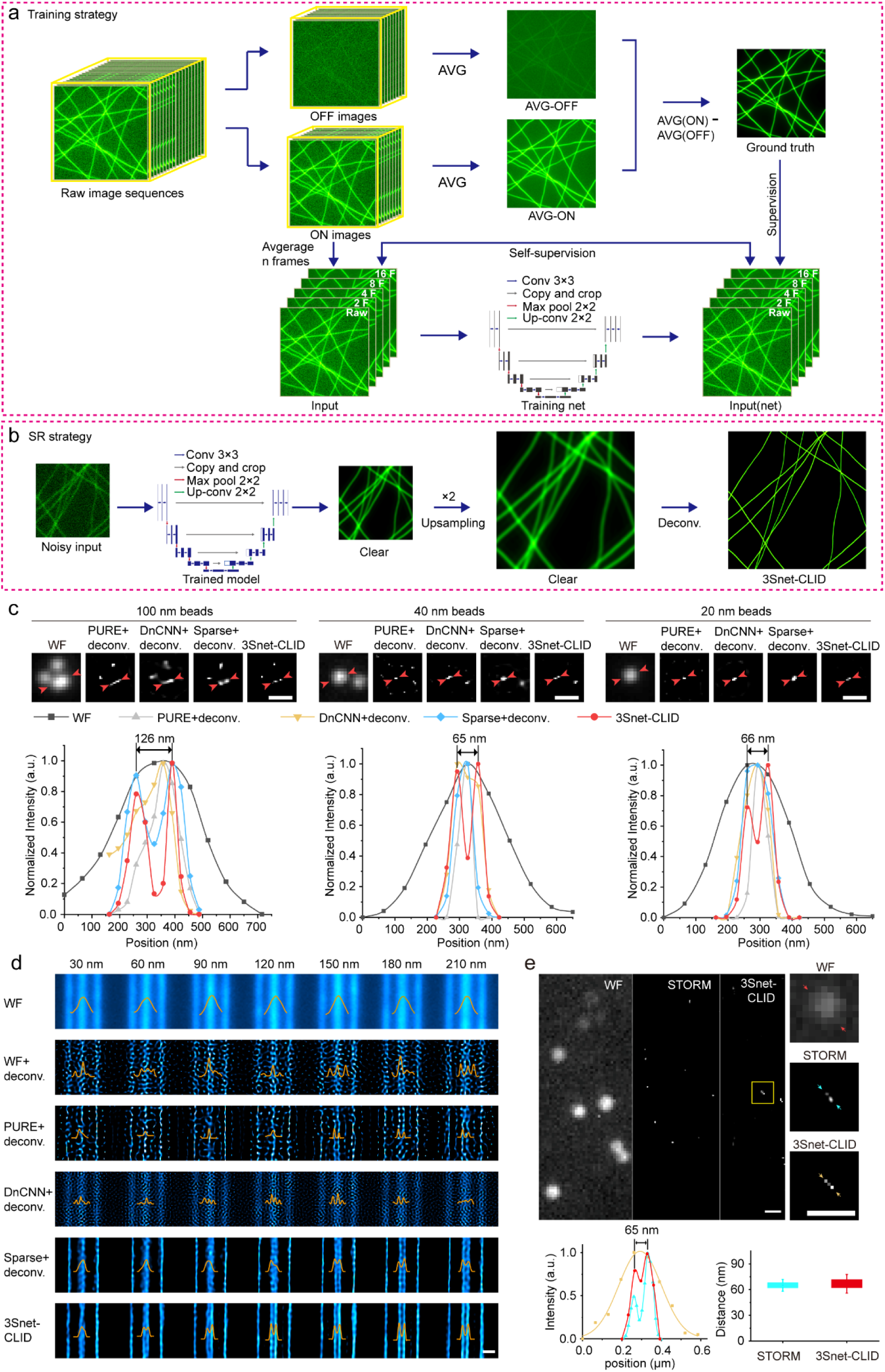
Workflow and validation of 3Snet-CLID using standard structures. **a**, Training strategy of 3Snet-CLID. The denoised clear images generated by 3S-denoising are utilized as ground truth for training supervision; Noisy images with different noise levels are obtained by averaging ON images with varying frames (1, 2, 4, 8, 16), and then two noisy images at each noise level are randomly selected as a pair to train the self-supervised network for every training iteration (**methods**). **b**, Trained 3Snet-CLID network. A single-frame noisy image is denoised, subsequently upsampled and applied to accomplish SR imaging. **c**, Two adjacent FluoSpheres beads with different sizes were effectively resolved using 3Snet-CLID. Top: Images of 100 nm, 40 nm and 20 nm beads under WF, PURE+deconvolution, DnCNN+ deconvolution, Sparse+deconvolution and 3Snet-CLID; Bottom: Intensity profiles along the lines indicated by the arrow heads in Top row head. The peak distances between two adjacent beads with different sizes resolved by 3Snet-CLID are 126, 65 and 66 nm, while full width at half maximum (FWHMs) of corresponding WF intensity profiles are 310, 254 and 251 nm, respectively. **d**, 3Snet-CLID has achieved a spatial resolution of 60 nm on the commercial Argo-SIM under wide field (WF) microscopy. The intensity profiles of the corresponding double-line pairs (30 nm, 60 nm, 90 nm, 120 nm, 150 nm, 180 nm and 210 nm distances) are displayed. **e**, Cross-validation of 3Snet-CLID and thunderSTORM. Left: DNA origami samples labeled with paired fluorophores of 60 nm distances imaged by WF, thunderSTORM and 3Snet-CLID; Middle: Zoomed-in regions of yellow box; Right: Intensity profiles along the lines indicated by the arrow heads and the average distance measured using two-peak gaussian fitting function (n = 7). Scale bar: 500 nm.

### 3Snet-CLID achieves 65-nm resolution of single frame WF image validated by standard known structures and cross-validation

We evaluated the performance of 3Snet-CLID in imaging structures with known ground truth obtained through WF microscopy. To do this, we compared the results of 3Snet-CLID, Sparse deconvolution, and direct deconvolution using other denoising methods, including Poisson unbiased risk estimate (PURE)^30^ and denoising convolutional neural networks (DnCNN)^31^, applied to the same image. Initially, we assessed ball-shaped fluorescent beads of varying sizes. Both 3Snet-CLID and Sparse deconvolution effectively resolved the 126-nm distance between two closely positioned 100-nm beads, whereas PURE and DnCNN deconvolution methods did not achieve this level of resolution (Fig. 2c). For smaller beads (20 and 40 nm), 3Snet-CLID succeeded in separating beads that were 65/66 nm apart— a feat that was unattainable with WF microscopy, Sparse deconvolution, PURE deconvolution, and DnCNN deconvolution (Fig. 2c). Following this, we utilized the linear structure of the commercial Argo-SIM as previously reported^18^. When lines were spaced at intervals of 60, 90, 120, 150, 180 and 210 nm, they were indistinguishable under WF, WF after RL deconvolution or DnCNN deconvolution but were clearly separated by 3Snet-CLID (Fig. 2d). Sparse deconvolution can resolve two lines separated by 150 nanometers (Fig. 2d), consistent with previous reports^18^. PURE deconvolution achieves the same spatial resolution as Sparse deconvolution but results in images with numerous artifacts (Fig. 2d). Subsequently, we conducted a cross-validation between 3Snet-CLID and STORM using DNA origami samples labeled with paired fluorophores. The origami structure incorporated two fluorescent sites labeled with FITC-488 molecules for repetitive optical selective exposure (STORM) microscopy, positioned 60 nm apart at the Nyquist sampling criteria edge (32 nm per pixel). While the paired molecules were hardly discernible under WF microscopy, they were well resolved by either 3Snet-CLID or STORM (Fig. 2e). Collectively, these findings highlight that 3Snet-CLID can effectively process structures of various shapes, achieving nearly a fourfold enhancement in spatial resolution (∼65 nm) compared to WF microscopy and nearly a twofold improvement compared to Sparse deconvolution.

### 3Snet-CLID enable live-cell single-frame SR imaging of cellular structures

We first assessed the imaging capabilities of 3S-CLID on single-frame biological samples from WF and SD imaging, using Sparse deconvolution for comparison. 3Snet-CLID significantly enhanced spatial resolution and contrast in WF imaging, allowing for clear visualization of actin meshes and filaments within a dense actin network in fixed HeLa cells. In contrast, images obtained using Sparse deconvolution lost many details of the actin structures. (Fig. 3a and 3b). Furthermore, 3Snet-CLID can be effectively utilized in dual-color imaging, improving resolution across different wavelengths to enhance visualization of fixed cellular structures (Fig. 3c). The dual-color imaging of F-actin (blue) and PMP (peroxisomal membrane protein, magenta) in fixed HeLa cells demonstrated that 3Snet-CLID produced sharper actin filaments with higher contrast compared to WF imaging (Fig. 3c). Additionally, it enabled the previously blurred fluorescent punctum of peroxisomes to be resolved as a ring-shaped structure located within the actin network. However, Sparse deconvolution was generally insufficient for resolving most of these ring-shaped structures, with the exception of particularly large rings. (Fig. 3c).

**Fig. 3.**
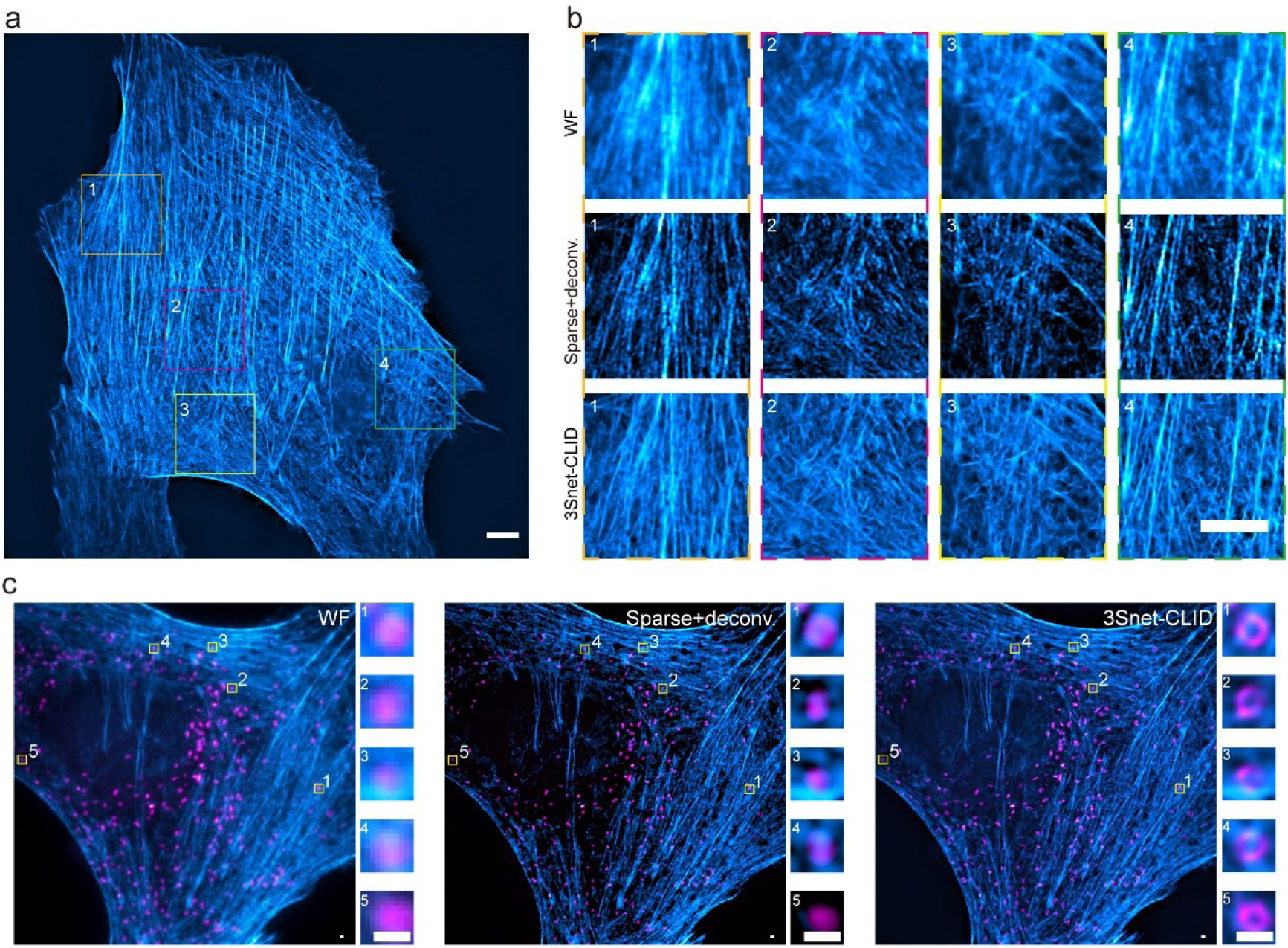
3Snet-CLID massively improves the quality of cellular structures. **a**, 3Snet-CLID imaging of F-actin labeled with phalloidin in fixed HeLa cells. **b**, Zoomed-in regions (1-4) from (**a**). Top to Bottom: WF, Sparse+deconvolution and 3Snet-CLID. **c**, Dual-color WF, Sparse+deconvolution and 3Snet-CLID imaging of F-actin with abberior STAR GREEN-labeled phalloidin (blue) and peroxisomes with abberior STAR RED-labeled PMP70 (magenta) in fixed HeLa cells. Colums: magnified views from white boxed regions. Scale bars: 5 μm (**a**, **b**), 500 nm (**c**).

In live Jurkat T cells, 3Snet-CLID enabled the analysis of dynamic actin changes during immune synapse formation with single-frame temporal resolution (Fig. 4, Supplementary Video 1). The enhanced spatial resolution facilitated detailed observation of the dynamic lamellipodial actin networks and lamellar actin networks corresponding to the distal supramolecular activation cluster (dSMAC) and peripheral supramolecular activation cluster (pSMAC), respectively (Fig. 4a). Both 3Snet-CLID and Sparse deconvolution techniques clearly resolved the dynamic retrograde flow of actin across the dSMAC and the centripetal movement of actin arcs across the pSMAC. Consistent with the previous report^32^, the rate of actin retrograde flow across the dSMAC appeared constant, as indicated by the linearity of the slopes in the kymographs for this region (Fig. 4a, dSMAC, red dash lines). However, contrary to the previous report^32^, we observed variable centripetal flow rates across the pSMAC, as reflected by different linearities in the kymograph slopes for this area (Fig. 4a, pSMAC, red dash lines). Furthermore, both 3Snet-CLID and Sparse deconvolution techniques clearly showed that some actin arcs at the pSMAC converge along the same trajectory over time (Fig. 4a, white arrow heads), highlighting the variation in movement speeds of these arcs. Unlike the uniform distribution seen with WF imaging, higher spatial resolution with 3Snet-CLID and Sparse deconvolution revealed that intensity distributions along the trajectories in both dSMAC and pSMAC are not uniform (Fig. 4a). This suggests possible dynamic assembly or disassembly of actin during movement. Correspondingly, the size of actin clusters at the outer edge of the pSMAC varies at different time points. In addition, consistent with the previous report^32^, the formation of concentric actin arcs at the boundary between the dSMAC and pSMAC remains unclear due to the resolution limits of WF imaging. Compared to Sparse deconvolution, 3Snet-CLID provides a clearer and easier observation of individual actin bundles or multiple bundles assembling into an actin arc at the boundary and moving inward across the pSMAC (Fig. 4a, yellow arrows).

**Fig. 4.**
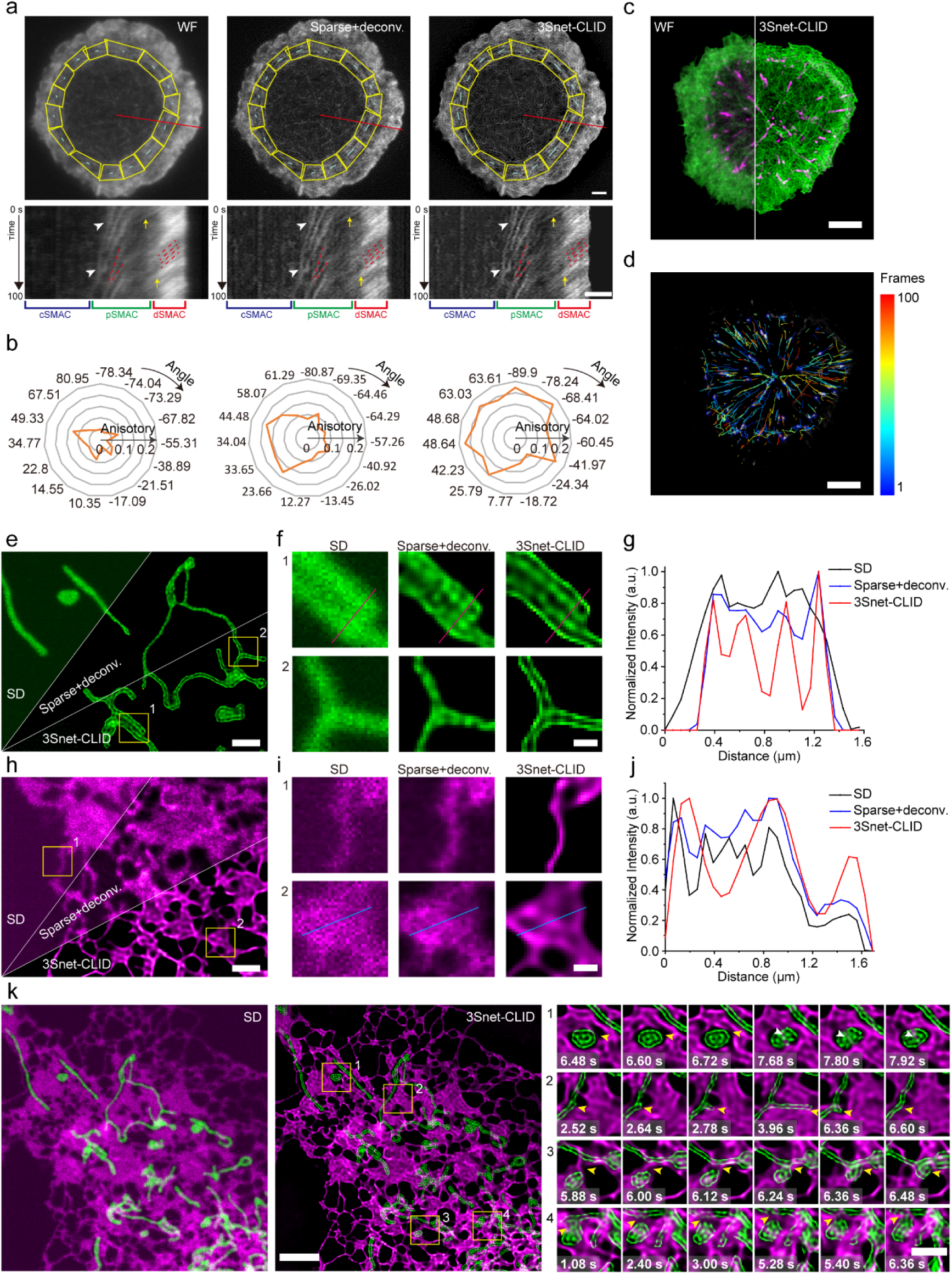
Single-frame live-cell SR imaging enabled by 3Snet-CLID. **a**, Utilizing 3Snet-CLID to analyze the intricate actin structures within the immune synapse of Jurkat T cells. Top: WF microscopy, Sparse-deconvolution and 3Snet-CLID imaging of F-actin labeled with F-tractin-StayGold in live Jurkat T cells. Bottom: Kymographs of F-tractin-StayGold in the region corresponding to the red line. **b**, The actin arcs anisotropy of the peripheral supramolecular activation cluster (pSMAC) in (**a**) measured by FibrilTool. Compared to WF microscopy and Sparse-deconvolution, 3Snet-CLID resolves more clear actin arc structure of the pSMAC, and provides more accurate quantified anisotropy data reflecting the effects of actin arc in the pSMAC organization. **c**, Dual-color imaging of F-tractin-StayGold-labeled F-actin (green) and EB1-mScarlet-I-labeled growing microtubule plus-ends (magenta) in living Jurkat T cells. **d**, Dynamic EB1 on growing microtubule plus-ends were automatically tracked over time using TrackMate. The color bar represents the frame number when different EB1 trajectories appear. **e**, SD, Sparse+deconvolution and 3Snet-CLID of TOM20-StayGold-labeled mitochrondria in live COS-7 cells. **f**, Zoom in regions (1, 2) from (**e**). **g**, Intensity profiles along the lines in top (**f**). **h**, SD, Sparse+deconvolution and 3Snet-CLID of Sec61β-mScarlet-I-labeled endoplasmic reticulum in live COS-7 cells. **i**, Zoom in regions (1, 2) from (**h**). **j**, Intensity profiles along the lines in bottom (**i**). **k**, Dual-color imaging of TOM20-StayGold-labeled mitochrondria (green) and Sec61β-mScarlet-I-labeled endoplasmic reticulum (magenta) in live COS-7 cells imaged by SD and 3Snet-CLID. Scale bars: 2 μm (**a**, **e**, **h**, **k** (right)), 5 μm (**c**, **d**, **k** (left)), 500 nm (**f**, **i**).

Studies have indicated that actin structures in different regions of the pSMAC layer exhibit diverse orientations and polarities, underscoring the importance of quantifying the anisotropy of these structures to understand their influence on pSMAC actin arc organization^33^. Through the analysis of angles and anisotropy radius using FibrilTool^34^, the actin arc anisotropy within the pSMAC region (Fig. 4a, highlighted in yellow and Fig. 4b) demonstrated that, due to enhanced resolution, 3Snet-CLID and Sparse deconvolution were more proficient in quantitatively assessing the orientation and polarity of actin arcs compared to WF microscopy. Additionally, compared to Sparse deconvolution, 3Snet-CLID produced greater actin arc anisotropy within the pSMAC region in various directions (Fig. 4b).

In a dual-color imaging mode, 3Snet-CLID enabled the visualization of dynamic movement of F-actin, labeled with F-tractin-StayGold (green) and microtubule plus-ends, labeled by EB1-mScarlet-I (magenta) (Fig. 4c, Supplementary Video 2). Employing a StarDist model provided in TrackMate^35^, microtubule plus-ends were automatically tracked over time (Fig. 4d). These tracking results demonstrated the ability of 3Snet-CLID to resolve the two poles of the microtubule-organizing center (MTOC) within the cell. Observations revealed the dynamic movement of EB1 proteins radially from the poles along the microtubule length towards the periphery, with varying densities of growing microtubules in different radial directions, indicating a polar generation and distribution of microtubules. Using single-molecule tracking technology, it was observed that the movement trajectories of EB1 molecules mostly did not overlap, with EB1 showing up at different growing microtubule plus ends at distinct time intervals. Interestingly, some EB1 molecules did not stem from the MTOC. As time passed, certain EB1 molecules disappeared earlier within the cell, while others vanished in the distant dSMAC layer of the immune synapse. These results suggest that during the immune synapse formation, EB1 molecules can detach and reattach from specific growing microtubules dynamically, indicating intermittent microtubule growth and dynamic circulation of EB1 proteins along the growing microtubules.

In a final evaluation, we examined the efficacy of dual-color 3Snet-CLID for live cellular structures using SD microscopy. At single-frame temporal resolution, 3Snet-CLID enabled visualization of the dynamic interactions between the endoplasmic reticulum (ER) and mitochondria (Supplementary Video 3). In the green channel, 3Snet-CLID clearly resolved the outer membranes of two closely spaced mitochondria that Sparse deconvolution could not distinguish, as well as the possibly stretched thin outer membranes of mitochondria (Fig. 4e-4g). Similarly, in the red channel, 3Snet-CLID demonstrated higher spatial resolution than Sparse deconvolution, allowing for the resolution of finer and sharper tubular ER structures, as well as tubulated ER sheet structures with meshes that Sparse deconvolution failed to resolve (Fig. 4h-4j). Notably, 3Snet-CLID effectively captured the dynamic changes in the outer membrane structure of mitochondria, a feature that SD cannot analyze, and accurately tracked the dynamic alterations in the tubular and sheet structures of the ER. Using 3Snet-CLID, numerous dynamic events were observed, such as the dynamic generation and disappearance of ER leading to mitochondrial morphology changes, as well as the stretching, recycling, fusion, and repetitive fusion-fission processes of mitochondria (Fig. 4k and Supplementary Video 3).

## Discussion

In our study, we introduced a novel denoising approach named synchronized signal switching (3S) denoising, which leverages optically modulated RSFP to improve image quality. By combining this denoising method with RL deconvolution, we introduced the 3S-CLID technique, enabling enhanced spatial resolution without the additional costly SR modules. Furthermore, the denoised images are utilized in training the deep learning network 3Snet, leading to the development of 3Snet-CLID for achieving single-frame SR imaging in live cells using standard FPs or dyes. Both 3S-CLID and 3Snet-CLID facilitate straightforward SR imaging using WF or SD microscopy, offering a seamless upgrade for existing microscopes without the need for instrumental modifications.

3S-CLID is an advanced SR imaging technique that integrates denoising with RL deconvolution. A key factor in the effectiveness of CLID is ensuring that the denoising process preserves signal integrity and maintains the distribution of signal across camera pixels, which is crucial for effective subsequent deconvolution. The impressive performance of 3S-CLID is largely due to its unique 3S denoising method. Unlike other denoising techniques that typically use intensity relationships between pixels and their neighbors (e.g., Hessian or Sparse methods^18, 36^, Gaussian filter^37^, wiener filter^38^) or remove high-frequency noise in the frequency domain (e.g., Wavelet transforms^39^, Low-pass filter^40^, the 3S denoising method employs label characteristics to extract useful signals and suppress noise, rather than relying solely on algorithmic noise reduction strategies. Traditional algorithmic denoising methods often alter the intensity distribution of the signal across pixels while removing noise. In contrast, the 3S denoising method processes each pixel independently based on the distinct modulation responses of signal and noise. This approach removes systematic noise through background subtraction within the same cell sample (ON-OFF) and combines image averaging to address Gaussian and Poisson noise. Consequently, it produces a denoised image that closely matches the noise-free ground truth, thereby enhancing the effectiveness of subsequent RL deconvolution and improving spatial resolution.

3Snet-CLID is an advanced DLSR imaging of living cells, providing high temporal and spatial resolution in WF and SD imaging. It enhances the resolution of single fluorescence images by nearly four-fold, achieving a spatial resolution of 65 nm. This is the highest resolution reported for such imaging without the need for an SR module. Compared to other DLSR technologies, it offers the following key features and advantages: 1) The 3Snet-denosing network combines supervised and self-supervised learning (Fig. 2a and 2b), with ground truth data improving the effectiveness of self-supervised learning, while self-supervised learning enhancing the robustness of supervised learning of new or underrepresented real images. 2) Obtaining the ground truth (clear image) is easy and simple. Remarkably, the clear images obtained with 3S denoising exhibit superior quality with reduced noise compared to those obtained through the conventional imaging averaging method, showcasing better preservation of signal integrity and distribution (Fig. 1b and 1c). This high-quality ground truth empowers the images generated by the 3Snet denoising network to enhance spatial resolution through direct RL deconvolution. 3) Using RSFP to acquire ground truth for training, the 3Snet denoising neural network effectively denoises samples labeled with conventional FPs and/or dyes, supporting 3Snet-CLID SR imaging. 4) Unlike analytical models that depend on assumptions about specimen and image properties to enhance resolution, 3Snet-CLID streamlines the process by removing the requirement for multiple parameter sets. This accessibility enables a diverse group of biologists to achieve reliable SR images consistently. 5) Compared to the the computation speed of Sparse deconvolution calculated according to the optimized parameters in the article^18^, the computation speed of 3Snet-CLID is 50 to 240 times of Sparse deconvolution (Supplementary Table 1). 6) In contrast to DLSR methods that use SR structures as ground truth^10, 41^, 3Snet-CLID provides two key benefits: Firstly, it eliminates the requirement to incorporate SR modules into WF or SD imaging systems and to use deep learning SR structures. Secondly, the denoising network of 3Snet-CLID has a simpler relationship between input and output data, with subsequent deconvolution being a separate process. As a result, the SR imaging effect is more robust and applicable to a wide range of cellular structures, such as lines, puncta, rings, tubules, and filaments, captured by WF and/or SD microscopes (Fig. 2).

Due to the widespread use and benefits of WF and confocal imaging, we expect that the versatility and effectiveness of parameter-free 3S/3Snet-CLID methods will lead to significant scientific advancements. Although our primary emphasis is on SR imaging in 2D WF and SD, these methods could potentially be applied in other WF-based techniques, like light sheet microscopy, and volumetric imaging to improve resolution.

## Methods

### CLID frameworks

For 3S-CLID, lifeact-SkylanS labeled F-actin in fixed U-2 OS cells were subjected to a series of acquisitions of ON and OFF frames. SkylanS were activated by 405 nm illumination (5 ms, 1.29 W/cm^2^) and multiple ON frames were captured under 488 nm illumination (5 ms, 5.39 W/cm²). 488 nm illumination was also used to switch off Skylan-S molecules. The same number of OFF frames were imaged with identical illumination settings as the ON frames. To obtain a clear image, we averaged the images captured in the ON state and subtracted the average from those taken in the OFF state. The denoised clear image were then upsampled by factor 2 using Lanczos. RL deconvolution was performed on the clear image to enhance its resolution, using Bessel function as the point spread function (PSF) following

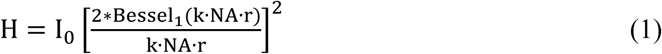

Where Bessel_1_ is the Bessel function of the first kind of order zero, I_0_ is the intensity of the incident radiation, k = 2*π*⁄*λ*, λ is the wavelength, NA is numerical aperture, r is the circular radius of the polar coordinate.

For 3Snet-CLID, the clear image obtained by 3S-CLID was used as the ground truth. In order to obtain the dataset for training 3Snet, we first extracted all ON state images from the raw ON/OFF image sequences. As shown in Extended Data Fig. 2, to generate training sets with diverse noise levels, we started by gathering 50 frames of images from the ON state and organizing them into five datasets using the combination formula *C*^*k*^, where n equals 50 and k is 1, 2, 4, 8 or 16. Next, we randomly selected 50 elements from each dataset and calculated the average of the k images in each chosen element. Thereafter we randomly selected two images in each dataset as the input (x) and target (y) images for the U-net model and using ground truth to supervise the training. The resulting predictions were calculated by the following loss function to execute the training stage.

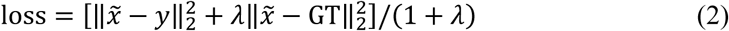

where *x̃* represents the network outcome of the input *x*, and 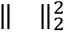 refers to the *l_2_* norm, λ is constraint weight of the loss function, GT is ground truth.

### Simulations of synthetic microtubule structures

To conduct benchmarks with simulated ground truth, microtubule-like structures were created using the ‘Random Walk’ process, and the maximal curvatures in the program were utilized^18^. The ground truth objects were then convolved with a microscope PSF (1.49 NA, 488 nm excitation, 65 nm pixel) to generate simulated blurred ground-truth images (GT_blur). Subsequently, the images were normalized to a 0-255 range and contaminated with Poisson noise (setting lam, the expected number of events occurring, to 5) and Gaussian noise (setting STD, the standard deviation of the distribution, to 5). The raw images were captured by a camera with a pixel size of 65 nm and pixel amplitudes of 16 bits.

### Optical setups

#### The wide field (WF) system

The WF system was based on a commercial inverted fluorescence microscope (IX83, Olympus) equipped with an objective (100×/NA 1.49 oil, UAPON, Olympus) and laser light with wavelengths of 405 nm (LBX-405-300-CIR-PP, Oxxius), 488 nm (Genesis MX488-500 STM, Coherent) and 561 nm (Jive 500-561 nm, Cobolt). Acoustic optical tunable filters (AOTFnC-400.650-TN, AA Opto-Electronic) were used to combine, switch, and adjust the illumination power of the lasers. A collimating lens (20×/NA 0.40, Plan N, Olympus) was used to couple the lasers to a polarization-maintaining single-mode fiber (QPMJ-3AF3S, Oz Optics).

The output lasers were collimated and then expanded by a beam expander (GCO-2503, Daheng Optics). The light passed through several mirrors (PF10-03-P01, Thorlabs) and a tube lens (ITL200, Thorlabs) was used to focus on the back focal plane of the objective. A multiband filter cube was used and emitted fluorescence collected by the same objective passed through a dichroic mirror (ZT405/488/561/640-phase R, Chroma) and an emission filter (FF01-446/523/600/647-25, Semrock). For dual-color imaging, an image splitter (Optosplit II, Cairn) was used before the sCMOS camera (Flash 4.0V2 C11440-22CU, Hamamatsu) to split the emitted fluorescence into two channels.

#### The Spinning-disk confocal microscopy

The Olympus SpinSR10 system utilized an inverted fluorescence microscope (IX83, Olympus) with four lasers (405 nm, 488 nm, 561 nm and 647 nm) and an objective (100×/NA 1.50 oil, Olympus). The setup included a scanning system (CSU-W2, Yokogawa) and a microscope incubator (Okolab) specifically designed for live cell imaging. The system is operated using CellSens Dimension software and the image capture is performed using an sCMOS camera (ORCA-Fusion BT, C15440-20UP, Hamamatsu).

#### The DeltaVision™ OMX microscope

The DeltaVision OMX system, developed by GE Healthcare, was equipped with an objective (60× PLAPON /1.42 NA), four lasers (405 nm, 488 nm, 561 nm, and 640 nm), four filter sets, and four sCMOS cameras, enabling high-speed imaging in 4-channel modes.

### Standard and cell samples for 3Snet-CLID imaging

#### DNA origami materials

The M13mp18 phage DNA sourced from New England BioLabs was used directly without additional purification. All staple strands, including Alexa Fluor 488-labeled and biotin-modified staple strands, were obtained from Tsingke Biotech (Beijing). The DNA origami staple strands were pre-mixed and stored at −20℃.

#### DNA origami design and assembly

The DNA origami nanostructure and synthesis was performed as previously described with slight modifications^42, 43^. Briefly, the DNA origami structures were assembled in a single reaction, including 10 nM M13mp18, 100 nM unmodified staple strands, 100 nM biotin-modified strands, and 400 nM Alexa Fluor 488-labeled strands with DNA-PAINT extensions in DNA origami folding buffer. The annealing process, carried out by a PCR device, entails initially holding the temperature at 25℃ for 20 seconds, then gradually increasing it to 90℃ over 15 minutes at a rate of 0.5℃ per second. Subsequently, maintaining the temperature at 90℃ for 1 minute, and then decreasing it by 1℃ per cycle at a rate of 0.1℃ per second until reaching 70℃. At 70℃, maintaining this temperature for 3 minutes, while decreasing the temperature to 30℃ by 1℃ per cycle at the same rate. After holding it for 30 seconds, the temperature is finally lowered to 4℃ at a rate of 0.5℃ per second.

For the purification process, a 100 kDa ultrafiltration concentrator was utilized. Initially, 500 μL of DNA origami purification buffer (15 mM MgCl_2_ in TAE buffer) was added and centrifuged at 5,000 g at 4°C for 7 min. Then, 50 μL of annealed solution was mixed with 450 μL of DNA purification buffer and centrifuged at 2,000 g at 4°C for 20 min. Subsequently, 450 μL of DNA purification buffer was added, and centrifugation at 2,000 g at 4°C for 20 min was repeated twice. The filter was then inverted into a fresh tube and centrifuged at 2,000 g at 4°C for 3 min, and the collected solution was stored at −20°C.

#### DNA-PAINT imaging

Confocal imaging dishes (Cellvis) were coated with 1 mg/mL biotin-labeled BSA (Solarbio) for 5 min at room temperature. After three washes with PBS, NeutrAvidin (1 mg/mL in PBS) was added to the microwell, incubated for 5 min, and washed three times with PBS. Subsequently, the 60 nm DNA origami structure was diluted in DNA origami imaging buffer (10 mM MgCl_2_ in PBS) and incubated for 5 min at room temperature, followed by three washes with the same buffer. Once the imager strands were added, the sample was ready for imaging. For the 60 nm structure, an exposure of 200 ms and an illumination power intensity of approximately 4-8 mW/cm^2^ were applied, with 100 frames acquired for the reconstruction. The acquired images were reconstructed with thunderSTORM the Fiji plugin^44^.

#### Argo-SIM slide

We also utilized a commercial fluorescent sample (the Argo-SIM slide, Argolight) with ground truth patterns consisting of fluorescing double line pairs (with spacing ranging from 0 nm to 390 nm; *λ_ex_* = 300-550 nm) to validate the resolution (http://argolight.com/products/argo-sim).

#### FluoSpheres imaging

To further validate the resolution, FluoSpheres of different sizes (20 nm-, 40 nm- and 60 nm-diameter, Thermo Fisher Scientific) were diluted with PBS and loaded onto glass coverslips. After allowing the fluorescent beads to settle for 30 min, a single-frame image was taken for analysis.

#### Live cell samples

For live 3Snet-CLID experiments, U-2OS or COS-7 cells were labeled with Lifeact-StayGold (Lifeact-SG)/Tom20-SG/Sec61-mScarlet-I using Lipofectamine™ 3000 transfection reagent (Thermo Fisher Scientific) according to the manufacturer’s instructions. Jurkat T cells, on the other hand, were transfected with F-tractin-SG/EB1-mScarlet-I via electroporation.

#### Fixed cell samples

For the 3S-CLID experiment, cells transfected with genetic indicators were fixed using 4% paraformaldehyde in PBS for 15 min at 37℃, followed by thorough washing with PBS.

#### Cell culture and preparation

The COS-7 and U-2OS cell lines were purchased from ATCC, while the Jurkat T cell line was acquired from the National Infrastructure of Cell Line Resources. COS-7 cells were cultured in DMEM (Gibco), and U-2OS cells were grown in McCoy’s 5A medium (Gibco), both supplemented with 10% FBS (Gibco), 100 U/mL penicillin, and 100 mg/mL streptomycin (Hyclone). Jurkat T cells were cultured in RPMI 1640 medium (Gibco) supplemented with 10% FBS, 1 mM sodium pyruvate solution, 50 μM β-mercaptoethanol (Gibco), and 0.1% Mycoplasma prevention reagent (Transgen) at 37℃ and 5% CO_2_. For imaging experiments, COS-7 or U-2OS cells were seeded onto glass coverslips, which were precoated with 10 μg/mL fibronectin for 30 min at 37℃. For imaging Jukat T cells, coverslips were coated overnight at 4°C with 2.5 μg/mL monoclonal anti-CD3 (eBiosciences) and washed with PBS three times. Cell imaging commenced within 10 min after loading cells onto coverslips.

#### Electroporation conditions

Cells were seeded at a density between 5×10^5^ and 1×10^6^ cells/mL to ensure that cells were in the exponential growth phase with high viability. Prior to electroporation, the cells were washed with PBS, counted, and resuspended in Opti-MEM™ medium (Gibco) to a cell density of 5 × 10^6^ cells/mL, and mixed with DNA plasmids (10 μg).

Following gentle mixing of the cells and plasmids in Opti-MEM™ medium (Gibco), the mixture was transferred into electroporation plates with a 0.2 cm gap (Bio-Rad). The electroporation process involved applying a voltage of 120 V, pulse interval of 0.1 s, 2 pulse cycles, and pulse lengths of 50 ms using square waveform pulses (Bio-Rad). Subsequently, 100 μL of the electroporated cells were transferred to 12-well plates containing 1 mL of culture media for Jurkat T cells and were then incubated at 37℃ for 24 hours.

### Date analysis

We calculated peak SNR (pSNR) values between reconstructed images (I) and ground truth (GT) images by:

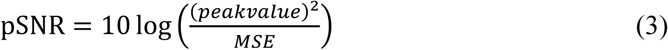

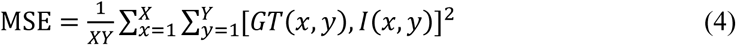

Structural similarity (SSIM) values between reconstructed images (I) and ground truth (GT) images by:

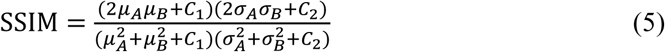

where *μ_A_* and *μ_B_* are the local means, *σ_A_* and *σ_B_* are the standard deviations for reconstructed images (A) and ground truth (B) images sequentially.

Standard deviation (STD) is obtained by:

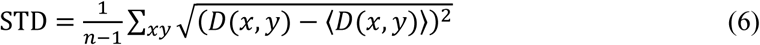

where n is the number of pixels; *D*(x, y) represents the difference between noisy images and ground truth images, and 〈*D*(*x*, *y*)〉 is the averaged intensity of the difference images.

To measure full width at half maximum (FWHM), the intensity profile is fitted by the normal distribution function as following:

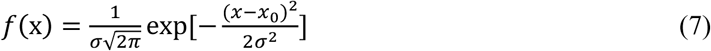

where *σ* is the standard deviation and *x*_0_ is the expected value, then FWHM is calculated by:

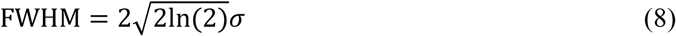

The anisotropy of actin arcs was examined using the Fiji plug-in FibrilTool^34^. FibrilTool was integrated into a macro that conducted orientation and polarity analysis within specified regions of interest nested within a grid-like layout that encompassed the overall image. The main fiber orientation was represented as a cyan line in each subregion of interest.

Object tracking was conducted utilizing the TrackMate^35^ plug-in for Fiji, and the maximum distance covered by EB1 tracks in each instance in Jurkat T cell was calculated using TrackMate as previously mentioned.

### Reporting summary

Further information on research design is available in the Nature Portfolio Reporting Summary linked to this article.

## Data availability

All the data that support the findings of this study are available from the corresponding author upon request.

## Code availability

The tutorials and the updating version of our 3Snet-CLID software can be found at https://github.com/FudongXue-xpyLab/CLID-py37.

## Acknowledgements

This project was supported by the National Natural Science Foundation of China (T2394513, 92254306, 21927813, 32227802 and 32027901), the National Key R&D Program of China (2022YFC3400600-2 and 2022ZD0211900), and the Strategic Priority Research Program of Chinese Academy of Sciences (XDB37040301). We thank Dr. Yuanyuan Li for the help with the DNA PAINT experiment (Institute of Biophysics, Chinese Academy of Sciences).

## Author contributions

P. X. and L. Y. conceptualized the project and provided detailed feedback and project direction. F. X., W. H., L. Y., Z. X., J. R., and C. S. implemented the strategy. F. X. and Z. X. ran analyses. P. X., L. Y., F. X. and W. H. wrote the paper.

---

**Extended Data Fig. 1.**
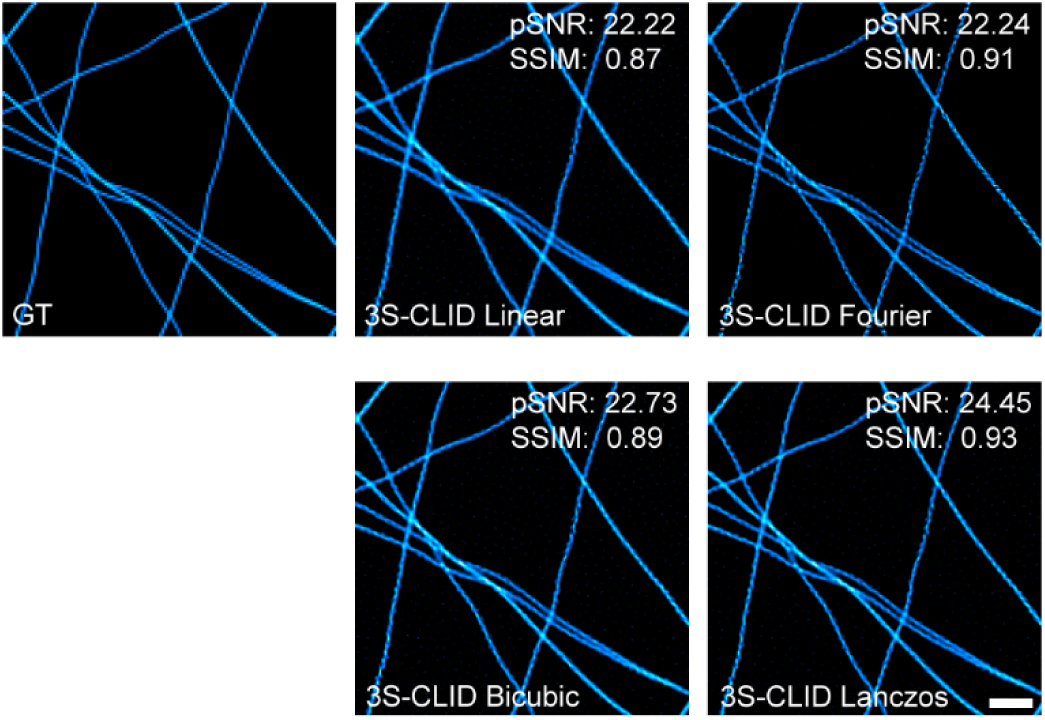
Assessment of different upsampling approaches for 3S-CLID. 3S-denoised synthetic structures were upsampled by linear, Fourier, Bicubic or Lanczos interporation approache for RL deconvolution. The pSNR and SSIM of 3S-CLID images were measured to compare the performance of the four upsampling approaches. Scale bar: 1 μm.

**Extended Data Fig. 2.**
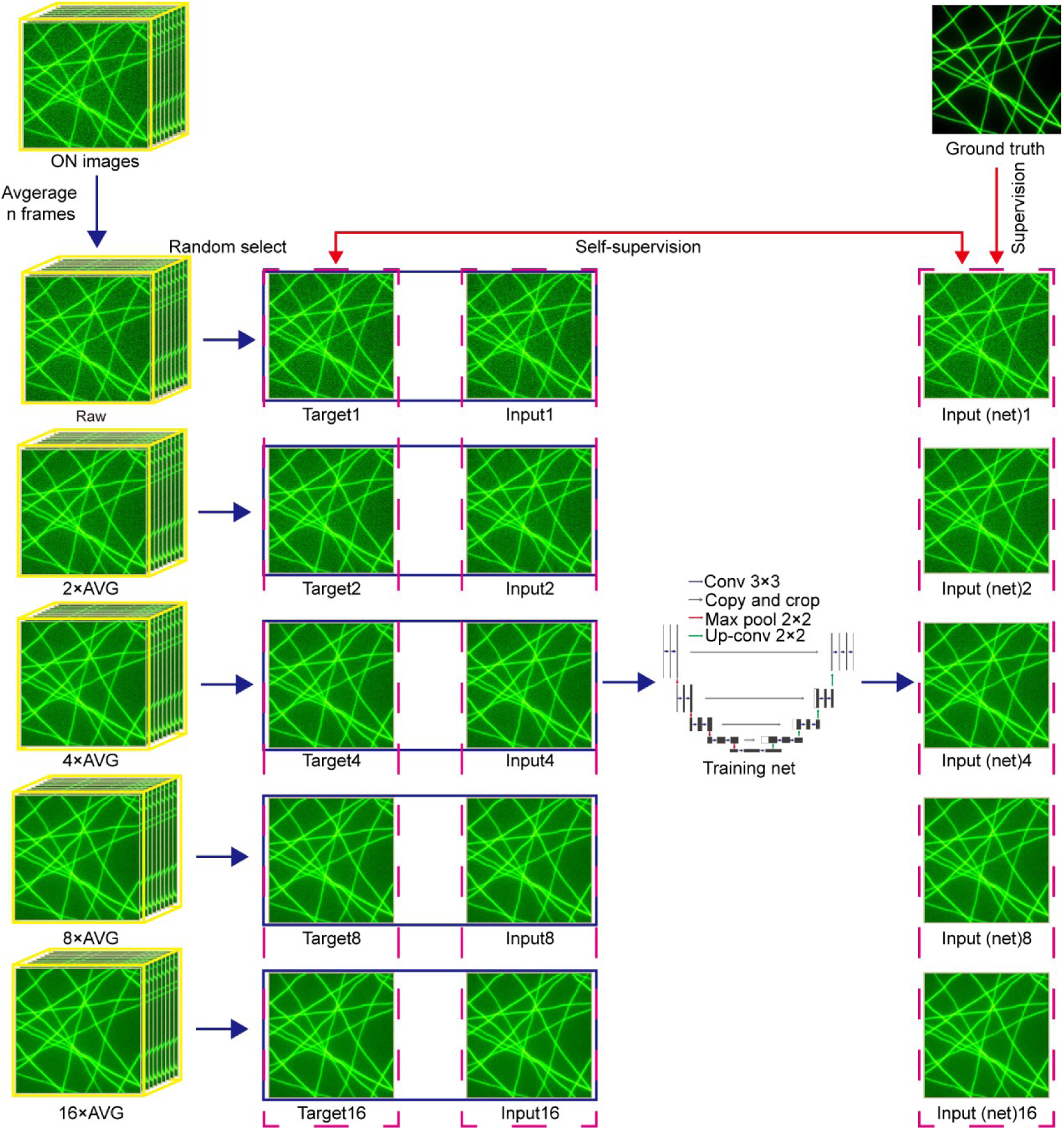
Dataset generation for 3Snet-CLID. To generate dataset for self-supervision, we first extract ON state images from the raw ON/OFF image sequences captured using SkylanS photoswitching. To generate training sets with diverse noise levels, 50 frames of images from the ON state are organized into five datasets using the combination formula *C*^*k*^, where n equals 50 and k is 1, 2, 4, 8 or 16. Next, 50 elements are randomly selected from each dataset and the average of the k images in each chosen element is calculated. Finally, two images in each batch are randomly selected as the input and target data to the U-net model. For supervision, the clear image obtained by 3S-denoising was used as the ground truth (**Methods**).

**Supplementary Table 1.** Running times of 3Snet-CLID and sparse-deconvolution for different figures.

|  | Pixels, No. of frames | Running time |  | Ratio of time<br>(Sparse/CLID) |
| --- | --- | --- | --- | --- |
|  |  | 3Snet-CLID | Sparse-deconv. |  |
| Fig.3b | 1024×1024, 1 | 9 s | 24 min | 160 |
| Fig.3c | 512×512, 1 | 5 s | 20 min | 240 |
| Fig.4a | 512×512, 100 | 1 min 37 s | 1 h 20 min | 49.5 |
| Fig.4e, h | 512×512, 100 | 3 min 13 s | 3 h 15 min | 60.6 |

